# Joint language production: an electrophysiological investigation of simulated lexical access on behalf of task partner

**DOI:** 10.1101/2020.09.18.303099

**Authors:** Anna K. Kuhlen, Rasha Abdel Rahman

**Affiliations:** Department of Psychology, Humboldt-Universität zu Berlin, Berlin, Germany

**Keywords:** language production, social interaction, semantic interference, electrophysiology, picture naming

## Abstract

This study investigates in a joint action setting a well-established effect in speech production, cumulative semantic interference, an increase in naming latencies when naming a series of semantically related pictures. In a joint action setting, two task partners take turns naming pictures. Previous work in this setting demonstrated that naming latencies increase not only with each semantically related picture speakers named themselves, but also with each picture named by the partner (Hoedemaker, Ernst, Meyer, & Belke, 2017; Kuhlen & Abdel Rahman, 2017). This suggests that speakers pursue lexical access on behalf of their partner. In two electrophysiological experiments (N=30 each) we investigated the neuro-cognitive signatures of such simulated lexical access. As expected, in both experiments speakers’ naming latency increased with successive naming instances within a given semantic category. Correspondingly, speakers’ EEG showed an increasing posterior positivity between 250-400ms, an ERP modulation typically associated with lexical access. However, unlike previous experiments, speakers were not influenced by their partner’s picture naming. Accordingly, we found no electrophysiological evidence of lexical access. To reconcile these findings we pooled behavioral data from five experiments (N=144). Within this large sample we find empirical evidence for partner-elicited interference. Furthermore, our data suggests that speakers may be less affected by their partner’s naming response in settings with remotely located task partners (as in present experiments). We conclude that speakers do not always represent their partner’s naming response and that our experimental setting may have limited the participants’ evaluation of the task as a joint action.

---

The intricate cognitive and neural processes underlying speaking have been investigated in copious tightly controlled experimental settings, in which single participants produce words or utterances (for review see e.g., Meyer, Roelofs, & Brehm, 2019). Recently, increasingly complex experimental settings have been introduced, in which two or more individuals interact and take turns speaking. This kind of setting approximates to a greater extent the setting in which people typically speak: together with conversational partners.

On the neural level, speaking to a conversational partner differs from speaking without communicative intentionality (Kuhlen, Bogler, Brennan, & Haynes, 2017; Willems et al., 2010). A series of additional demands are placed upon the human cognitive system when speaking in a conversational setting (Hagoort & Levinson, 2014). One of these demands is the close temporal coordination found between speakers and listeners. Conversational partners in natural dialogue alternate between speaking and listening at a remarkable speed (Stivers et al., 2009). It has been proposed that this type of close coordination is achieved by predicting how the partner’s incoming utterance may end, enabling speakers to plan their response in advance (e.g., Bögels, Magyari, & Levinson, 2015). These predictions are assumed to rely on representations similar to those engaged during speaking (Pickering & Garrod, 2013). The present study aims to contribute to an understanding of the neurocognitive processes engaged when two speakers alternate speaking by comparing electrophysiological brain activity recorded when speakers speak themselves with activity recorded when their task partner speaks.

Recent behavioral evidence suggests that speakers engage in lexical access for words that they anticipate their task partner to say. For example, we have investigated speech production in a setting in which two task partners take turns naming pictures of everyday objects. This setting is based on the continuous naming task, in which pictures are presented in a seemingly random order. Embedded in this stream of pictures are pictures that are semantically related to each other, for example, a series of different kinds of birds.

In the classic setting, in which only one speaker names these kinds of pictures it is a robust finding that with each additional picture speakers name within a given semantic category their naming response becomes slower (roughly by 20ms), so-called cumulative semantic interference (see e.g., Belke, 2013; Costa et al., 2009; Howard et al., 2006; Navarrete et al., 2010). One prominent explanation for this effect is lexical competition between semantically related lexical entries: when retrieving representations associated with a particular lexical entry semantically related entries are co-activated and compete with the target entry for selection. Once the target lexical entry is selected, the connection, either between the target concept and its lexical entry (Howard et al., 2006) or between the concept and its semantic features (Belke, 2013), is strengthened, and therefore previously named lexical entries become stronger competitors when co-activated during the subsequent retrieval of a semantically related item. Alternative accounts of this effect propose that connections to non-target lexical entries are weakened and subsequently take longer to be selected (e.g., Oppenheim, Dell, & Schwartz, 2010; Navarrete et al., 2010).

In our previous set of experiments (Kuhlen & Abdel Rahman, 2017) we were able to demonstrate that semantic interference can be elicited not only when speakers themselves name semantically related pictures, but also when the pictures are named by the task partner. Speakers’ naming latencies increased more steeply within a given semantic category when the task partner named additional within-category pictures compared to when these additional pictures were presented only visually. Presumably this partner-elicited interference arises because those pictures named by the partner trigger lexical access and thus contribute to the number of strong lexical competitors activated during subsequent speech production. This supports the proposal that speakers activate, during their partner’s speaking, representations similar to those activated when naming pictures themselves.

We observed partner-elicited interference not only when speakers could hear their task partner name pictures (Kuhlen & Abdel Rahman, 2017, Experiment 1), but also when speakers could not hear the partner name the pictures either because their partner was located in a different room (Kuhlen & Abdel Rahman, 2017, Experiment 2) or because they were wearing noise-canceling headphones (Kuhlen & Abdel Rahman, 2017, Experiment 3). Thus only the belief that the partner was naming a picture seemed to be sufficient to engage lexical representations on behalf of the partner.

The finding that speakers engage in lexical access for words assigned to the task partner is corroborated by a study from Hoedemaker and colleagues (2017), who investigated how pictures named by a task partner influence subsequent speech production. Applying a comparable joint picture-naming task this study demonstrated that a speaker’s naming latencies increased as a function of the number of semantically related pictures that had previously been named by the partner. The authors conclude that speaking and listening rely on shared representations.

Applying a different type of picture-naming paradigm, a study by Baus and colleagues (2014) provides first electrophysiological evidence that speakers engage in lexical processing in anticipation of their task partner speaking. This study investigated event-related potentials (ERPs) recorded while the task partner named pictures of high or low lexical frequency and compared them to potentials observed when these pictures were presented only visually. A positive inflection around 320-420ms after picture onset that was most pronounced at posterior electrode sites was modulated by lexical frequency. A similar posterior positivity at an earlier latency has been interpreted as an index of lexicalization (cf. Costa et al., 2009). Most notably, this component was observed in trials in which the task partner named pictures, but not in trials in which pictures were presented only visually. A similar posterior positivity, starting somewhat earlier, was observed when participants themselves produced low-frequency versus high frequency words. The observation of similar components both while participants named pictures and while partners named pictures supports the proposal that speakers simulate their partner’s speech production (Pickering & Garrod, 2013). In addition, this study reported a second positivity from 420 to 550ms in those trials in which the partner named a picture. An increased positivity in this time window has also been observed in nonverbal tasks when contrasting trials in which a task partner acts with trials in which nobody acts (Demiral, Gambi, Nieuwland, & Pickering, 2016; Sebanz, Knoblich, Prinz, & Wascher, 2006; Tsai, Kuo, Hung, & Tzeng, 2008; Tsai, Kuo, Jing, Hung, & Tzeng, 2006). It has been interpreted to reflect inhibitory processes engaged when representations related to action preparation need to be suppressed. Taken together there is converging evidence, both from behavioral and electrophysiological studies, for the assumption that speakers engage in (and possibly later inhibit) lexical processing on behalf of a task partner.

Yet, empirical evidence is not unanimous: some studies report that speakers do not appear to represent their task partners’ utterances to the degree that they seek lexical access on behalf of their partner. A study by Gambi et al. (2015) comes to the conclusion that speakers represent the fact that their partner speaks, but finds no evidence that the speaker represents what the partner says (i.e. does not engage in lexical access). In this study speakers named pictures concurrently with a task partner who was located remotely in another room. Speakers were instructed that simultaneous to their own picture naming a second speaker was either naming the same picture or a different picture, didn’t name the picture, or performed a semantic categorization task. Speakers were able to name the pictures faster when the other speaker was presumably not naming pictures or was performing the categorization task. However, crucially, naming latency was not affected by the belief that the other speaker was naming the same or a different picture. If speakers had represented the second speaker’s action to the degree that they pursued lexical access, the authors reasoned, naming should have slowed down when speakers believed partners to be naming different pictures compared to naming the same pictures. These conclusions were corroborated by a recent study (Brehm, Taschenberger, & Meyer, 2019), which applied a similar task setting. In this study participants’ naming latencies were also affected by the partner’s task (yet, interestingly, in the opposite direction found in Gambi et al., 2015). But participants were not consistently affected by whether the partner concurrently named the same or a different picture. Both studies concluded that speakers do not represent in detail what their partner is saying.

A study by Hoedemaker and Meyer (2019) comes to a similar conclusion: In this study sets of three pictures were presented simultaneously on a display shared between a speaker and a task partner. Speakers always named the first picture on the display. The authors compared trials in which also the second picture on the display was named by speakers with trials in which the second picture was named by the task partner or by nobody. With the help of eye trackers speakers’ gaze shift from the first to the second picture was analyzed as a proxy for whether and when speakers prepared for naming the second picture. It was reasoned that if a partner’s picture naming elicits processes comparable to those when naming the picture oneself, gaze patterns in the trials in which speakers name the pictures should be comparable to those trials in which the partner names the pictures. Indeed, speakers looked at the second pictures more often and earlier when their task partner named it compared to when nobody named it. But, crucially, when speakers had to name pictures themselves they looked at the second picture significantly earlier and longer. An analysis of the speakers’ naming latencies suggests that speakers may not have represented their partner’s naming at all: speakers were slower to initiate the naming of the first picture when they also had to name the second picture. However, they were equally fast in initiating naming when the second picture was named by the partner or by nobody. The authors conclude that speakers engage in a more complete representation of pictures they name themselves and partially represent pictures named by the task partner.

In sum, evidence about speakers engaging in lexical processing on behalf of a task partner’s speaking is mixed. Some studies demonstrate that the semantic relatedness or word frequency of the pictures named by the partner affect speakers’ own language processing. These effects are hard to explain without assuming that speakers engage in lexical access. Yet other studies come to the conclusion that speakers do not, or only partially, represent their partner’s utterances and do not engage in lexicalization. Present study

With the current set of experiments we aim to contribute to our understanding of the processes underlying language production in social interaction by investigating the electrophysiological underpinnings of partner-elicited cumulative semantic interference. Specifically, we investigate whether speakers anticipate and simulate the content of a task partner’s naming in a variant of the continuous naming task in which two task partners take turns naming pictures. As a proxy for lexical processing we resort to an ERP modulation that has been associated with semantic interference. This modulation, a relative positivity associated with related compared to unrelated semantic contexts around 250-400ms at posterior electrode sites, aligns with the time course of lexical access. And its amplitude increases with additional category members named in the continuous naming task (Costa et al., 2009; Rose & Abdel Rahman, 2016) and with semantically related compared to unrelated distractors in the picture word interference task (Rose et al., 2018). Due to its tuning to lexical processing this component has been interpreted to reflect processes of lexical access (for critical discussion see e.g., Indefrey, 2011; Piai, Riès, & Knight, 2015; Strijkers & Costa, 2016).

On the basis of our previous work we employ a joint picture naming setting in which two task partners take turns naming a series of pictures showing everyday objects (Kuhlen & Abdel Rahman, 2017). The pictures can be grouped into different semantic categories of 10 exemplars each. Five exemplars were named by the participants; five additional exemplars were either named by the partner (Joint Naming condition) or were presented only visually (Single Naming condition). Thus, in both conditions participants named an equal number of semantically related pictures; what differed was whether, interspersed, additional pictures of the same category were named by the partner, or whether they were presented visually but were not named by anyone.

During the picture naming participant and task partner were seated in separate rooms (as Kuhlen & Abdel Rahman, 2017, Experiment 2). Thus, participants did not receive auditory feedback from their partner’s naming response. The shared task setting was defined by their belief of performing the task together with their remote partner, as well as by the minimal feedback they received from the speed of their partner’s supposed naming response (for details see below). The advantage of this setting is that our main contrast, trials in which task partners named pictures and trials in which pictures were presented only visually, were maximally identical to each other on a perceptual level (i.e. in both cases participants did not have to respond and they also did not hear anyone respond) and the EEG recordings were kept free from processes associated with speech perception.

Based on previous studies we expected an increase in naming latencies with each member of a semantic category named by the participant, i.e. ordinal position (cumulative semantic interference, see e.g., Belke, 2013; Costa et al., 2009; Howard et al., 2006; Navarrete et al., 2010). In addition we expected, based on previous studies in a joint action setting, that naming response slows due to the task partner naming semantically related pictures (partner-elicited semantic interference; Hoedemaker et al., 2017; Kuhlen & Abdel Rahman, 2017). To further understand the processes behind these effects we measured participants’ EEG. In line with previous work, we expected to observe an increased positivity with an onset around 250ms after stimulus onset at posterior electrode sites in response to the degree of interference experienced when naming semantically related pictures (Costa et al., 2009; Rose & Abdel Rahman, 2016). That is, we expected an increase in amplitude with each category member named by the participants. Crucially, we hypothesized that this marker can be observed not only when participants themselves speak, but also when their task partner speaks. We would interpret this as indicating that participants are simulating lexical access on behalf of their task partner.

The materials, experimental procedures, and analysis approach were kept identical to Experiment 2 reported in Kuhlen and Abdel Rahman (2017), with the exception that we aimed to increase statistical power for the EEG study by increasing the number of participants (from 24 to 30 participants) and the number of times stimuli were presented (from one time to three times). The current study was pre-registered prior to initiating data collection (Kuhlen & Rahman, 2017). Because Experiment 1 did not yield the expected findings, we sought to improve the basic experimental setting in Experiment 2. Both experiments were approved by the local Ethics Committee at the Humboldt-Universität zu Berlin.

## Experiment 1

### Methods

#### Participants

Based on the effect sizes observed in our previous experiment (Kuhlen & Abdel Rahman, 2017, Experiment 2) we simulated the outcome of the anticipated linear mixed model with 1000 iterations (simr package, Green, MacLeod, & Nakagawa, 2016). In this model the predictor naming condition was contrast coded using the sliding difference contrast (Joint naming: −0.5, Single Naming: 0.5), and the predictor naming order was centered and entered as continuous variable. In the simulation we predicted log-transformed naming latencies and entered as estimates .02 for the predictor naming order, −.01 for the predictor naming condition, and −.01 for the interaction between these two predictors. With 30 participants we reached a power estimate of 83% chance (95% confidence interval: 80.53, 85.28) for detecting the hypothesized partner-elicited cumulative semantic interference, namely, an interaction between naming order (cumulative semantic interference) and naming condition (Joint Naming vs. Single Naming).

We recruited healthy, right-handed, native German-speaking volunteers between the ages 18-36. Participants were reimbursed or received credit towards their curriculum requirements. Data from six participants were replaced because they expressed considerable doubt during the debriefing that their task partner had indeed been naming the pictures assigned to them. Data from two additional participants had to be replaced because of excessive trial loss due to noisy EEG recordings. The final data set consisted of 21 women and nine men. All participants gave informed consent prior to participating in the study.

#### Materials

Three hundred and twenty colored pictures (photographs) of man-made or natural objects were used as stimulus material (identical to those used in Kuhlen & Abdel Rahman, 2017). The objects map onto 32 different semantic categories (e.g., birds, beverages, flowers). Each category consisted of 10 exemplars. As Filler items served 120 additional objects, which were unrelated to the categories underlying the target items. All pictures were scaled to 3.5cm x 3.5cm and were presented on a homogenous grey background.

Pictures were collated in stimulus lists, which were created for each participant individually with the following constraints: The order in which pictures of one category were presented was randomly selected (by the program “Mix”, van Casteren & Davis, 2006). Pictures of one category were separated randomly by a minimum of two and a maximum of six unrelated pictures (separating pictures could be fillers or pictures belonging to different categories). To avoid a conceptual merging of two or more related categories (e.g., categories fish and birds merging to the superordinate category animal) these never overlapped within a list (for comparable procedure see Kuhlen & Abdel Rahman, 2017; Rose & Abdel Rahman, 2016).

The assignment of categories to naming condition (Joint vs. Single Naming) was counterbalanced across participants. In both naming conditions, half of the exemplars of a given category were assigned to the participant (participant names object). Under the Joint Naming condition, the other half of exemplars were assigned to the partner (partner names object); under the Single Naming condition, the other half was assigned to visual presentation only (nobody names object). The assignment of pictures to trials in which participants named the picture and trials in which the partner or nobody named the picture was random with the following two exceptions: trials, in which the participant speaks, were separated by maximally three trials, in which the participant does not speak. The first and the last presented exemplars of a category were always assigned to the participant. All filler items were also assigned to the participant.

Each participant was presented with the complete set of pictures three times (three repetitions). Pictures assigned to participants in the first round were also assigned to participants in the following repetitions. The same picture never appeared at the same ordinal position.

#### Procedure

Two participants were invited to the lab at one time. They were introduced to each other as task partners. Participants were told they would jointly name pictures while seated in different rooms (“similar to a virtual computer game”). Participants were instructed that pictures would be presented one at a time. A colored frame around the picture indicated who was to name the picture: the participant, the partner, or nobody. Participants were instructed to name as fast and accurately as possible those pictures coded in the assigned color. In all other trials they were told to do nothing. The corresponding color codes were assigned randomly at the beginning of each experiment.

After receiving the task instructions, participants were separated and seated in their respective experimental cabins for mounting the EEG cap. During this time, participants had the opportunity to study the pictures (presented unsorted on paper) and their corresponding name. Once both EEG caps were successfully mounted the two task partners were familiarized with the trial structure by 15 practice trials (using pictures not included in the main experiment). After the practice trials, experimenters coordinated the start of the main experimental session. Unbeknownst to the participants, the experimental sessions were run independently from each other.

Each trial began with a fixation cross of 500ms. The picture was then presented for a maximum duration of 2000ms. Pictures disappeared once the voice key was triggered by speech onset. In trials in which the task partner presumably named the picture, pictures disappeared at a randomly chosen interval corresponding to naming latencies observed in a past experiment (analog to procedures in Experiment 2, Kuhlen & Abdel Rahman, 2017). In trials in which nobody named the pictures, the pictures disappeared at a fixed interval of 2000ms. A blank screen of 1500ms followed each picture presentation and then the next trial began.

Each time participants and their supposed task partners completed naming the entire set of pictures (i.e., after each repetition) they took a break during which we disconnected them from the amplifier and asked them to step out of the cabin into the main room where they could socialize with their presumed task partner.

After the picture-naming task was completed, participants were separately interviewed on their beliefs about the experiment. In particular, the experimenters assessed whether participants believed they had truly been working together with their partner on the same task. If the participants expressed substantial doubt that they had named pictures together with their task partner (indicated as 7 or higher on a ten-point scale) their data were excluded from further analyses.

### Data acquisition and analyses

#### Naming response

Naming latencies (reaction times) were recorded with the help of a voice-key, which marked the time that elapsed between the onset of the picture and the onset of the naming response. The experimenter marked false alarms of the voice-key (e.g., due to speech disfluency) as well as other erroneous trials (object named wrongly or by the wrong person). Analyzed were naming latencies for all valid target trials in which participants named the picture. Trials were marked as invalid if participants failed to produce a naming response, if the naming response was wrong or disfluent or if the voice key was erroneously triggered. This concerned 6% of all target trials. The remaining valid trials were corrected for outliers, excluding all trials with naming latencies below 200ms or above two standard deviations from the individual’s overall mean value, 5% of all valid trials. In total 89.2% of all target trials was analyzed. Naming latencies were log-transformed prior to entering the Linear Mixed Model (see below).

Electrophysiology. Continuous EEG was recorded during the main experimental session with 62 Ag/AgCl electrodes arranged according to the extended 10/20 system. An electrode over the left mastoid was used as reference. The sampling rate was 500 Hz. Eye movements and blinks were registered by an external electrode near the lateral canthus of the left eye. Electrode impedance was kept below 5kOhm. Offline the EEG was re-referenced using the average reference transformation and low-pass filtered (high cutoff = 30 Hz, 24 db/oct). Eye movement and blink artifacts were removed through the software program Brain Electrical Source Analysis (BESA), which derives an estimate of the spatial distributions of ocular artifacts to be corrected from the EEG. Afterwards, EEG data were segmented in epochs of 2100ms, starting 100ms before the onset of the target (baseline interval). Remaining artifacts were addressed by automatic artifact rejection and by excluding segments with potentials exceeding 50μV and a threshold of 200μV. Finally, segments were averaged for each ordinal position within each naming condition.

EEG recorded during or before picture-naming may be contaminated by artifacts associated with speaking. The EEG was therefore corrected by Residue Iteration Decomposition (RIDE), a procedure that decomposes ERPs into component clusters with distinct trial-to-trial variabilities (in our case: stimulus-locked, response-locked, and clusters of variable latency). Due to the distinct characteristics of these components the procedure is well suited for identifying and eliminating articulation artifacts during overt speech production (Ouyang et al., 2016; see also Rose, Aristei, Melinger, & Abdel Rahman, 2019), see Figure 1.

**Figure 1.**
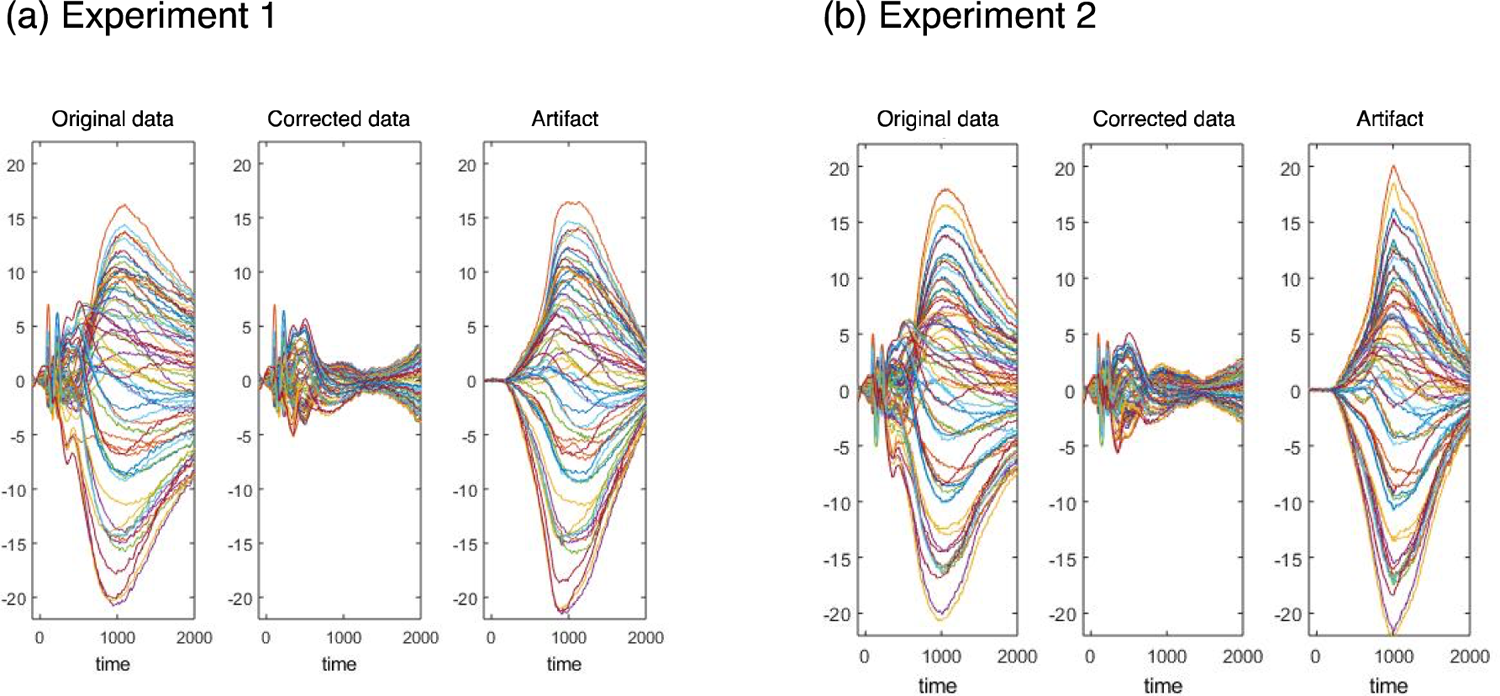
Decomposition of ERP data into component clusters based on residue iteration algorithm (RIDE) applied to trials in which participants name pictures for (a) Experiment 1 and (b) Experiment 2. Within each panel are displayed grand average ERP waveform before RIDE (left), stimulus locked component (middle) and response-locked component (right). See online article for the color version of this figure.

Analyzed were EEG data obtained from trials in which participants named the picture, trials in which the partner named the picture, and trials in which the picture was presented only visually. Based on previous reports of ERP effects in the continuous naming paradigm (cf. Costa et al., 2009; Rose & Abdel Rahman, 2016) we focused our analysis on mean amplitude modulations at posterior electrode sites at a time window typically associated with lexical access (Indefrey & Levelt, 2004; Indefrey, 2011). Specifically, our time window, 250-400ms after picture onset, as well as our region of interest (namely, Cp3, Cp4, P5, P3, Pz, P4, P6, PO3, POz, and PO4) was identical to those reported in Rose and Abdel Rahman (2016).

#### Analyses

Linear mixed effects models (LMM; Baayen, Davidson, & Bates, 2008) as implemented in the lmer function of the lme4 package (Bates, Mächler, Bolker, & Walker, 2014) for R (R Development Core Team, 2012) with random intercepts for participants and semantic categories^1^ were applied to our dependent measures, log-transformed naming latencies and single-trial EEG recordings averaged over our target time window and electrodes.

Our dependent measures were modeled as a function of the predictors naming condition (Joint Naming vs. Single Naming), naming order (ordinal position 1 to 5), and repetition (1 to 3). The predictors naming condition and repetition were contrast coded using the sliding difference contrast, which compares average latencies between neighboring positions. The predictor naming order was centered and entered as continuous variable.

Models were initially run with a maximum random effects structure (cf. Barr, Levy, Scheepers, & Tily, 2013). Using singular value decomposition, the initial full random effect structure was simplified, if necessary, by successively removing those random effects for which variance was estimated to be zero until the maximal informative model was identified.

For fixed effects, we report fixed effect estimates, standard errors (SE), and t values, as well as the estimates of variance and the square root (standard deviations) of the random effect structure, and goodness-of-fit statistics. Fixed effects were considered significant if |t| ≥ 1.96 (cf. Baayen, Davidson, & Bates, 2008).

As announced in our preregistration (Kuhlen & Abdel Rahman, 2017) cluster-based permutation tests as implemented in FieldTrip (Maris and Oostenveld, 2007) were performed on all electrodes on a time window between 100-700ms for exploratory data analyses (cf. Frömer, Maier, & Abdel Rahman, 2018). The permutation distribution was based on 1000 iterations, and considered clusters significant at an alpha-level of p < .05.

## Results

### Naming latencies

Participants took on average 876ms to name a picture (Joint Naming 877ms; Single Naming 876ms). Within a given semantic category, participants’ naming latencies increased on average by 10ms with each additional picture named, demonstrating cumulative semantic interference. The increase observed in the current experiment was somewhat smaller compared to our previous experiment in the same setting, in which we observed an increase of 17ms (Kuhlen & Abdel Rahman, 2017, Experiment 2).

Other than expected, pictures named by the task partner did not appear to elicit semantic interference (no partner-elicited semantic interference), as indicated by a comparable average increase in naming latencies in both naming conditions (Joint Naming: 9ms; Single Naming: 11ms); see Figure 2a. The LMM analyses confirmed these effects as follows, for summary see Table 1.

**Figure 2.**
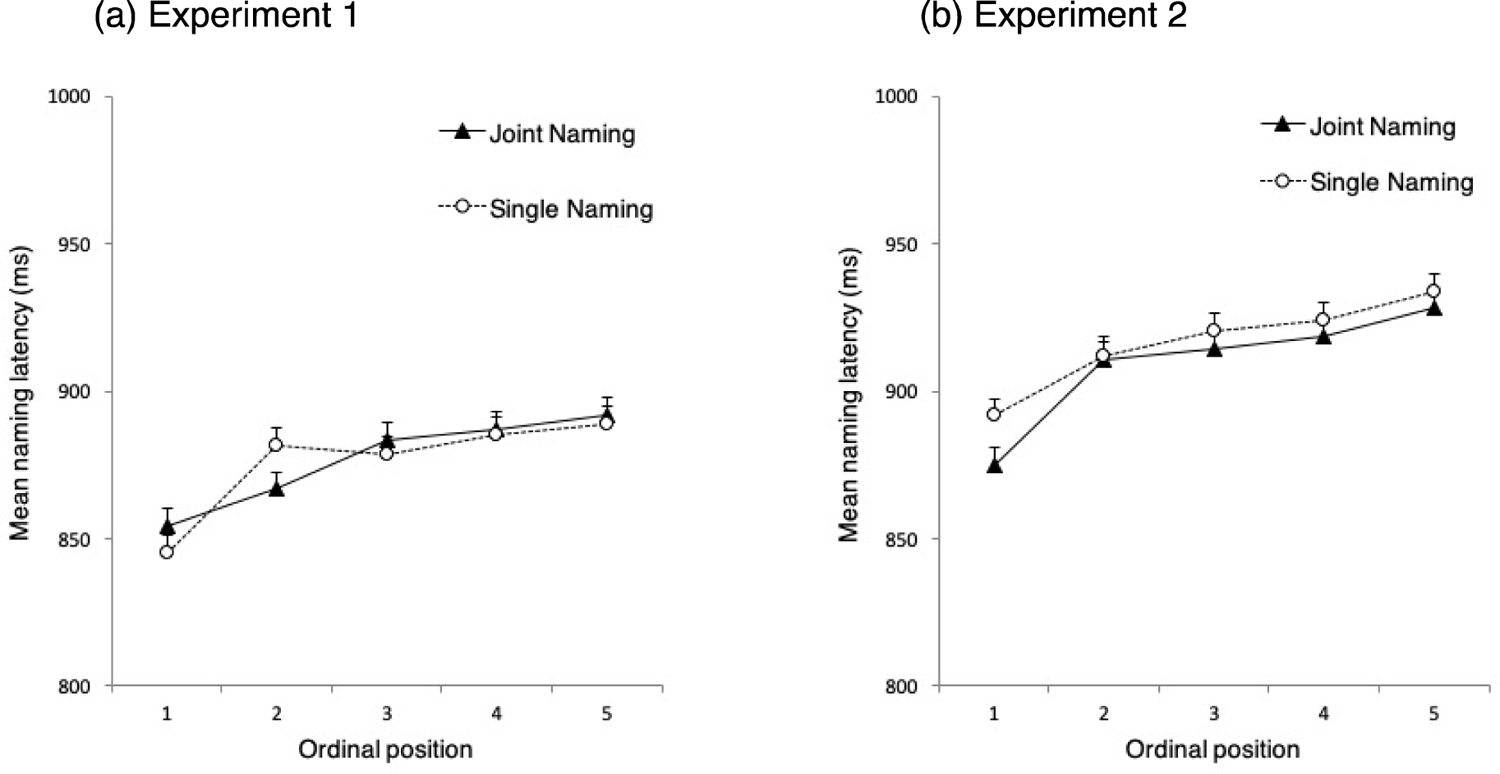
Mean naming latency and standard error (in milliseconds) for each ordinal position and naming condition. (a) Results of Experiment 1. (b) Results of Experiment 2.

**Table 1:**
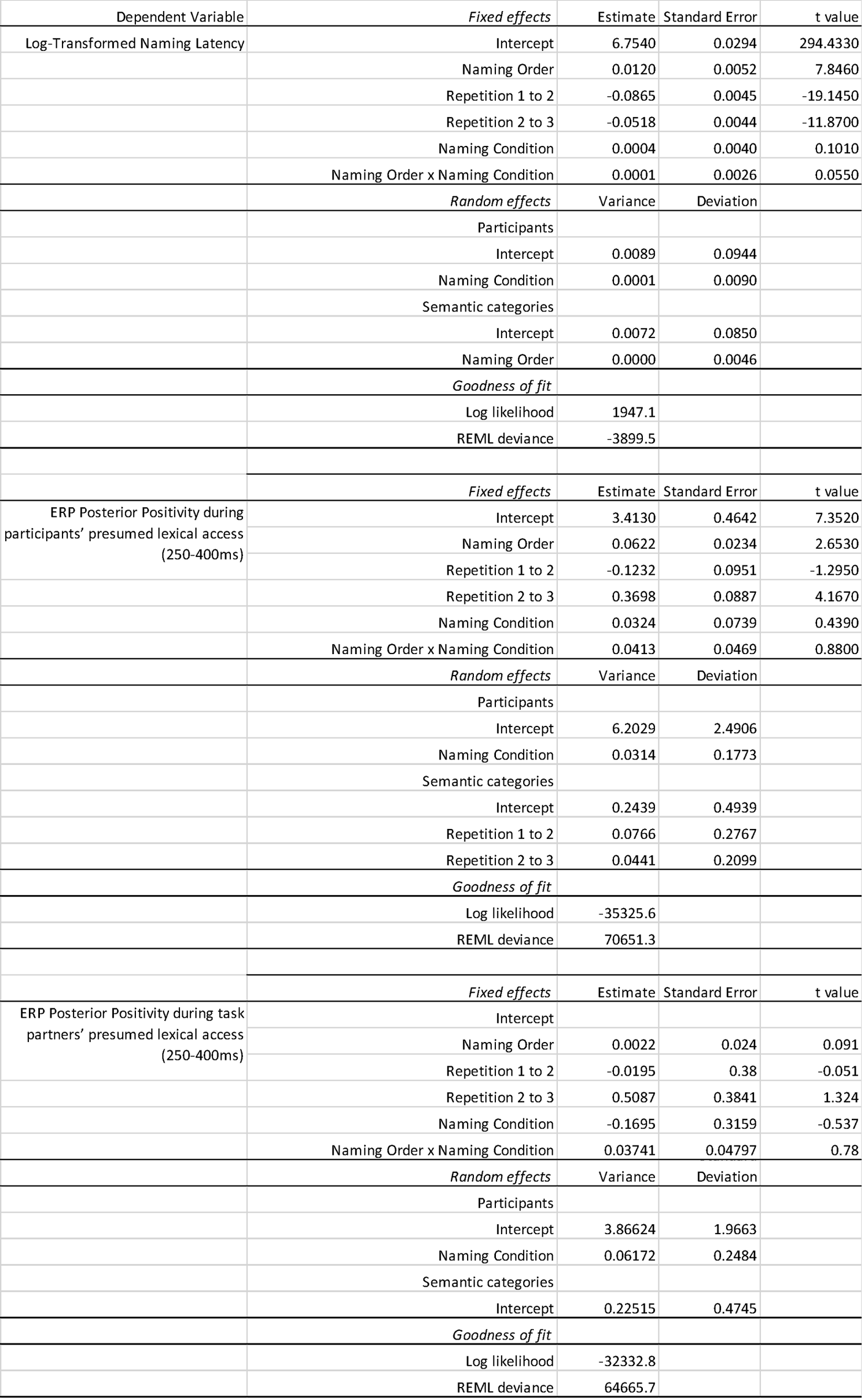
Fixed-effect estimates, standard errors, and t-values for the selected models of Experiment 1; estimates of the variance and square root (standard deviations) of the random effect structure and goodness-of-fit statistics. Fixed effects are considered significant if |t| ≥ 1.96 (cf. Baayen, Davidson, & Bates, 2008).

| Dependent Variable | Fixed effects | Estimate | Standard Error | t value |
| --- | --- | --- | --- | --- |
| Log-Transformed Naming Latency | Intercept | 6.7540 | 0.0294 | 294.4330 |
|  | Naming Order | 0.0120 | 0.0052 | 7.8460 |
|  | Repetition 1 to 2 | -0.0865 | 0.0045 | -19.1450 |
|  | Repetition 2 to 3 | -0.0518 | 0.0044 | -11.8700 |
|  | Naming Condition | 0.0004 | 0.0040 | 0.1010 |
|  | Naming Order x Naming Condition | 0.0001 | 0.0026 | 0.0550 |
|  | *Random effects* | Variance | Deviation |  |
|  | Participants |  |  |  |
|  | Intercept | 0.0089 | 0.0944 |  |
|  | Naming Condition | 0.0001 | 0.0090 |  |
|  | Semantic categories |  |  |  |
|  | Intercept | 0.0072 | 0.0850 |  |
|  | Naming Order | 0.0000 | 0.0046 |  |
|  | *Goodness of fit* |  |  |  |
|  | Log likelihood | 1947.1 |  |  |
|  | REML deviance | -3899.5 |  |  |
| ERP Posterior Positivity during participants' presumed lexical access (250-400ms) | *Fixed effects* | Estimate | Standard Error | t value |
|  | Intercept | 3.4130 | 0.4642 | 7.3520 |
|  | Naming Order | 0.0622 | 0.0234 | 2.6530 |
|  | Repetition 1 to 2 | -0.1232 | 0.0951 | -1.2950 |
|  | Repetition 2 to 3 | 0.3698 | 0.0887 | 4.1670 |
|  | Naming Condition | 0.0324 | 0.0739 | 0.4390 |
|  | Naming Order x Naming Condition | 0.0413 | 0.0469 | 0.8800 |
|  | *Random effects* | Variance | Deviation |  |
|  | Participants |  |  |  |
|  | Intercept | 6.2029 | 2.4906 |  |
|  | Naming Condition | 0.0314 | 0.1773 |  |
|  | Semantic categories |  |  |  |
|  | Intercept | 0.2439 | 0.4939 |  |
|  | Repetition 1 to 2 | 0.0766 | 0.2767 |  |
|  | Repetition 2 to 3 | 0.0441 | 0.2099 |  |
|  | *Goodness of fit* |  |  |  |
|  | Log likelihood | -35325.6 |  |  |
|  | REML deviance | 70651.3 |  |  |
| ERP Posterior Positivity during task partners' presumed lexical access (250-400ms) | *Fixed effects* | Estimate | Standard Error | t value |
|  | Intercept |  |  |  |
|  | Naming Order | 0.0022 | 0.024 | 0.091 |
|  | Repetition 1 to 2 | -0.0195 | 0.38 | -0.051 |
|  | Repetition 2 to 3 | 0.5087 | 0.3841 | 1.324 |
|  | Naming Condition | -0.1695 | 0.3159 | -0.537 |
|  | Naming Order x Naming Condition | 0.03741 | 0.04797 | 0.78 |
|  | *Random effects* | Variance | Deviation |  |
|  | Participants |  |  |  |
|  | Intercept | 3.86624 | 1.9663 |  |
|  | Naming Condition | 0.06172 | 0.2484 |  |
|  | Semantic categories |  |  |  |
|  | Intercept | 0.22515 | 0.4745 |  |
|  | *Goodness of fit* |  |  |  |
|  | Log likelihood | -32332.8 |  |  |
|  | REML deviance | 64665.7 |  |  |

The maximal informative model included random slopes for the factor naming condition for participants and random slopes for the factor naming order for semantic categories. This model showed a significant main effect for the predictor naming order, confirming cumulative semantic interference. The predictor repetition also yielded a main effect, indicating that naming latencies decreased from the first to the second and from the second to the third repetition, as can be expected due to practice effects. Most relevant for the current study, the hypothesized interaction between naming order and naming condition was not significant, providing no support for partner-elicited semantic interference.

Electrophysiological response while participant names pictures. As expected, we found a posterior positivity between 250-400ms after picture onset at posterior electrode sites. This positivity increased in amplitude from the first to the last picture named within a given semantic category, see Figure 3a. We interpret this modulation to reflect the processes underlying cumulative semantic interference (cf. Costa et al., 2009; Rose & Abdel Rahman, 2016). The increase of this positivity within a given semantic category did not differ in significant ways between the Joint Naming and Single Naming conditions. This corresponds with our behavioral findings.

**Figure 3.**
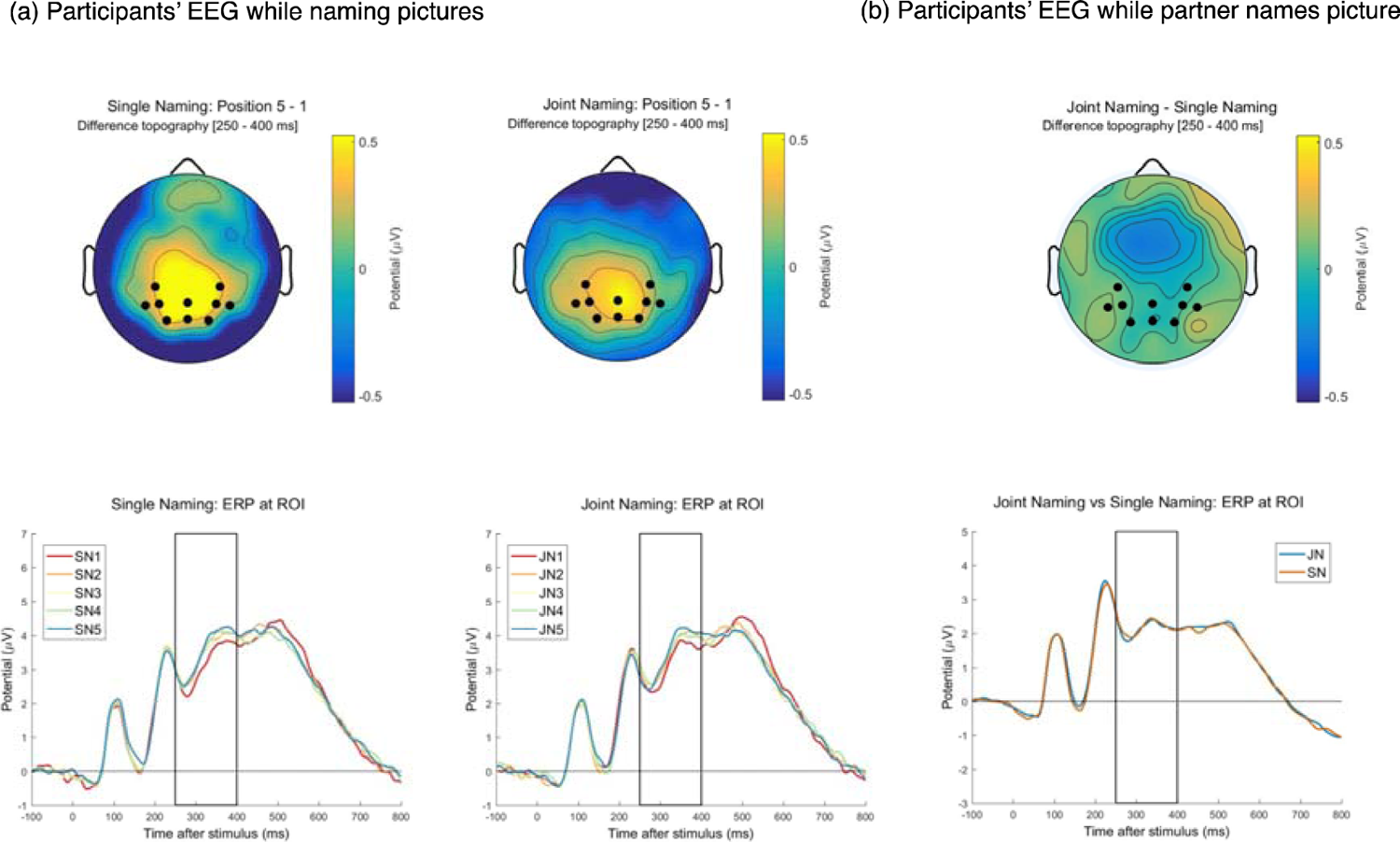
EEG results from Experiment 1. (a) Topographies (top) and waveforms (bottom) of event-related potentials elicited during participants’ presumed lexical access (250-400ms) when naming pictures of the fifth relative to the first ordinal position within a given semantic category for Single Naming (SN; right) and Joint Naming (JN; left) separately. (b) Topographies (top) and waveforms (bottom) of event-related potentials elicited in participants during trials in which they believe their partner is naming pictures (Joint Naming; JN) relative to trials in which pictures are presented only visually (Single Naming; SN) in the time-window of presumed lexical access (250-400ms). The electrodes of the region of interest are marked in black in the topography and were pooled to generate the waveforms. The time window of interest is marked by black box in the waveform. See the online article for the color version of this figure.

Our LMM analyses supported these observations, see Table 1. Our selected model contained random slopes for naming condition to model effects by participants and random slopes for repetition to model effects by semantic categories. This model showed a significant main effect for the predictor naming order, confirming on an electrophysiological level cumulative semantic interference. There was also a main effect for the predictor repetition, indicating that our target ERP modulation differed from repetition 2 to 3. This reflects a greater positivity at posterior electrode sites in the third repetition. Since this effect was not replicated in Experiment 2 we will not discuss it further.

Cluster based permutation tests were used to explore further our data beyond the defined region and time window of interest. In particular, previous studies (Costa et al., 2009; Rose & Abdel Rahman, 2016) have reported a relative negativity at posterior electrode sites starting around 400ms after picture onset. In one study this modulation decreased in amplitude (i.e., more negative) with ordinal position (Rose & Abdel Rahman, 2016).

To test for an effect of naming order we compared the first to the last picture named within a given semantic category. As expected, this analysis confirmed the posterior positivity reported in the LMM analyses above. The corresponding cluster was significant (p = .00) in a time window of approximately 275-425ms, covered our region of interest and reached beyond it into central electrode sites (for details, see Figure A, Panel a in Appendix A). Furthermore, we found a significant negative cluster (p = .01) in posterior electrode sites in the time window of approximately 500-600ms, see Figure A, Panel b in Appendix A. The time window and electrode positions overlaps with the negative-going modulation reported in Rose and Abdel Rahman (2016), and has been associated with the N400. Lastly, we found a second significant positive cluster (p = .00) starting approximately at 490ms and reaching up until our cutoff of 700ms after picture onset, covering frontal and central electrode sites, see Figure A, Panel a in Appendix A.

In a second permutation test we contrasted the Joint Naming with the Single Naming condition to test for an effect of Naming Condition. No significant clusters were identified.

Furthermore, a third permutation test investigated the interaction between naming condition and naming order by comparing the difference between the first and last picture naming within a semantic category for Joint Naming and Single Naming respectively. This analysis also did not yield significant clusters.

Electrophysiological response during partners’ lexical access. We searched in participants’ EEG for evidence of simulated lexical access in form of a comparable posterior positivity in those trials in which task partners named pictures.

Contrasting EEG recordings during the naming of the first and the last picture of a given semantic category, the topographies in the time window 250-400ms (same as applied in analysis of trials in which participants named the picture) did not give any visual evidence of such a posterior positivity for either naming condition, Joint Naming and Single Naming. Accordingly, there was also no evidence for a positivity when directly contrasting trials in which the task partner named the picture (Joint Naming) with those trials in which nobody named the picture (Single Naming), see Figure 3b. The corresponding LMM found no significant contribution of our predictors Naming Order, Naming Condition and Repetition. Please refer to Table 1 for model details.

Previous literature has reported specific components in the EEG related to processes involved in inhibiting a response. One such component, often called No-Go P300, is reduced in No Go relative to Go trials. This component may be larger for those trials that require the task-partner to respond compared to those that nobody responds to (for comparable findings Demiral, Gambi, Nieuwland, & Pickering, 2016; Sebanz et al., 2006; Tsai et al., 2008, 2006). Additionally, an early, negative modulation, peaking around 200ms after stimulus onset (N2), has been shown to characterize No Go trials, and may be more pronounced when a (simulated) task partner’s response needs to be inhibited.

We employed cluster-based permutation tests across all electrodes in search of a difference between our two types of No-Go trials, Joint Naming and Single Naming. This analysis yielded no significant clusters differentiating the two conditions. We view this as further indication that our participants did not represent their partner’s actions.

## Discussion

Consistent with the literature, this experiment demonstrates that speakers experience interference when naming semantically related pictures. This interference cumulates in size with each additional picture named within a given semantic category. Cumulative semantic interference was evidenced in participants’ behavior, as an increase in naming latency, and in participants’ brain response, as an increase in posterior positivity while participants were preparing their naming response (250-400ms after picture onset). Furthermore, we found evidence for an increased posterior negativity 500-600ms after picture onset. These observations replicate quite precisely previous studies on cumulative semantic interference (Costa et al., 2009; Rose & Abdel Rahman, 2016).

Yet the main goal of this experiment was to investigate the neurocognitive processes underlying partner-elicited cumulative semantic interference. Unfortunately, we were not able to replicate previous findings that a task partner’s picture naming creates additional interference (Hoedemaker et al., 2017; Kuhlen & Abdel Rahman, 2017). In the current experiment, speakers’ naming latencies were not affected by their task partner’s prior naming of semantically related pictures. Correspondingly, there was also no stronger increase in posterior positivity in the Joint Naming compared to the Single Naming condition when participants named pictures, and, crucially, also no sign of lexicalization in participants’ EEG while the presumed task partner was naming a picture. Thus, participants in the current experiment did not seem to engage in lexical access on behalf of their task partner.

Previous studies in joint action settings have suggested that task partners inhibit their tendency to execute a response on the task partner’s behalf (e.g., Sebanz et al., 2006; Tsai et al., 2008, 2006). If such inhibitory processes are engaged early enough they could effectively occlude or prevent lexicalization on behalf of a task partner. However, our exploratory analyses do not give any indication of greater inhibitory responses in trials in which task partners named pictures compared to trials in which nobody named pictures.

We therefore propose that the most parsimonious explanation for our data pattern is that participants did not represent their partners’ naming response. Although we have successfully demonstrated partner-elicited interference in an almost identical setting (Kuhlen & Abdel Rahman, 2017, Experiment 2), we speculate that the physical separation of the two task partners discouraged participants from perceiving the task as a joint activity. Indeed, many of our participants indicated in our debriefing interview that they did not experience the experimental setting as a joint action (German term: “Zusammenarbeit”). Possibly, because of the rather long time period during which the EEG cap was mounted (about 45-60 minutes), and during which the two task partners were separated from each other, our participants had simply forgotten about their task partner once they started the naming session. And during the naming session there was little evidence to remind participants of their task partner’s involvement.

Nevertheless, the physical separation of the two task partners was instrumental to exclude an influence of the obviously confounding factor that comes with a setting in which partners are physically co-present: namely that participants would hear the presented picture being lexicalized by the task partner in the Joint Naming condition while in the Single Naming condition the picture would be presented only visually with no auditory lexicalization. In Experiment 2 we therefore opted to keep the experimental control a remotely located task partner permits while strengthening the participants’ belief in the involvement of their remotely located task partner.

## Experiment 2

### Methods

#### Participants

We recruited healthy, right-handed, native German-speaking volunteers between the ages 18-36 who had not taken part in Experiment 1. Comparable to Experiment 1, the final data set consisted of 30 participants (19 women). Data from five participants had to be replaced because they doubted their task partner had been naming pictures during the partner-assigned trials. Data from two additional participants had to be replaced because of excessive trial loss due to noisy EEG recordings. Participants were reimbursed or received credit towards their curriculum requirements. All participants gave informed consent prior to their participation.

#### Materials

The materials were identical to those used in Experiment 1.

#### Procedure

The procedures followed in large parts the procedures of Experiment 1, with the following exceptions. After the two participants were introduced to each other and had received an initial overview over the joint naming task they were separated for mounting the EEG caps. After both EEG caps were mounted the two participants were brought together into one cabin where they underwent a practice session consisting of 15 trials, sitting side-by-side in front of the computer screen. After the practice trials ended, the participants were separated once again. The two experimenters pretended to coordinate the start of the experiment across the two cabins (while in reality the two experimental sessions started independent from each other). As in Experiment 1, participants were encouraged to socialize with each other during the two breaks.

Because some participants of Experiment 1 had indicated that the naming latencies of their supposed task partners had not appeared to be realistic, we sought to improve our mock naming latencies by assigning each stimulus picture with the typical (mean) naming latency we observed for this particular picture across four previous experiments (Experiment 1-3 reported in Kuhlen & Abdel Rahman, 2017, and Experiment 1 of current study).

Naming latencies, electroencephalographic recordings, and data analyses. The recording of participants’ naming latencies and EEG followed identical protocols as in Experiment 1. Of all target trials 6% were excluded due to a voice key error or wrong, disfluent or missing naming responses. Of the remaining trials 5% were excluded based on our outlier correction (cf. Experiment 1). In total, 88.99% of all target trials was analyzed.

As in Experiment 1, log-transformed naming latencies and mean amplitude modulations at our posterior region and time window of interest were modeled as a function of the predictors naming condition (Joint Naming vs. Single Naming), ordinal position (1 to 5), and repetition (1 to 3) in Linear Mixed Models. Cluster permutation tests were performed for exploratory analyses.

## Results

### Naming latencies

Participants took on average 913ms to name a picture (Joint Naming 909ms; Single Naming 916ms). Comparable to Experiment 1, naming latencies increased on average by 12ms with each additional picture named within a given semantic category, giving further evidence for cumulative semantic interference. Yet also in Experiment 2 the pictures named by the task partner did not elicit semantic interference: naming latencies increased at roughly the same rate in the Joint Naming and the Single Naming condition (Joint Naming 13ms; Single Naming 10ms), see Figure 2b. These conclusions were supported by our LMM analyses, for summary see Table 2.

**Table 2:** Fixed-effect estimates, standard errors, and t-values for the selected models of Experiment 2; estimates of the variance and square root (standard deviations) of the random effect structure and goodness-of-fit statistics. Fixed effects are considered significant if |t| ≥ 1.96 (cf. Baayen, Davidson, & Bates, 2008).

| Dependent Variable | Fixed effects | Estimate | Standard Error | t value |
| --- | --- | --- | --- | --- |
| Log-Transformed Naming Latency | Intercept | 6.794 | 0.024 | 282.978 |
|  | Naming Order | 0.1218 | 0.0015 | 8.227 |
|  | Repetition 1 to 2 | -0.0558 | 0.0045 | -12.418 |
|  | Repetition 2 to 3 | -0.0454 | 0.0044 | -10.409 |
|  | Naming Condition | 0.006 | 0.0038 | 1.564 |
|  | Naming Order x Naming Condition | -0.002 | 0.0026 | -0.769 |
|  | *Random effects* | Variance | Deviation |  |
|  | Participants |  |  |  |
|  | Intercept | 0.0110 | 0.1056 |  |
|  | Naming Condition | 0.0000 | 0.0070 |  |
|  | Semantic categories |  |  |  |
|  | Intercept | 0.0064 | 0.0802 |  |
|  | Naming Order | 0.0000 | 0.0042 |  |
|  | *Goodness of fit* |  |  |  |
|  | Log likelihood | 2002.2 |  |  |
|  | REML deviance | -4004.5 |  |  |
|  | *Fixed effects* | Estimate | Standard Error | t value |
| ERP Posterior Positivity during participants' presumed lexical access (250-400ms) | Intercept | 2.5000 | 0.4684 | 5.3360 |
|  | Naming Order | 0.1011 | 0.0255 | 3.972 |
|  | Repetition 1 to 2 | -0.0466 | 0.0880 | -0.5290 |
|  | Repetition 2 to 3 | -1.0649 | 0.0869 | -0.1900 |
|  | Naming Condition | 0.0647 | 0.0799 | 0.8090 |
|  | Naming Order x Naming Condition | -0.0072 | 0.0509 | -0.1510 |
|  | *Random effects* | Variance | Deviation |  |
|  | Participants |  |  |  |
|  | Intercept | 6.3144 | 2.5129 |  |
|  | Naming Condition | 0.0375 | 0.1938 |  |
|  | Semantic categories |  |  |  |
|  | Intercept | 0.2447 | 0.4947 |  |
|  | *Goodness of fit* |  |  |  |
|  | Log likelihood | -36369.6 |  |  |
|  | REML deviance | 72739.3 |  |  |
|  | *Fixed effects* | Estimate | Standard Error | t value |
| ERP Posterior Positivity during task partners' presumed lexical access (250-400ms) | Intercept | 1.195 | 0.4644 | 2.574 |
|  | Naming Order | 0.0009 | 0.0261 | 0.038 |
|  | Repetition 1 to 2 | -0.3277 | 0.4148 | -0.79 |
|  | Repetition 2 to 3 | 0.0147 | 0.4145 | 0.035 |
|  | Naming Condition | -0.496 | 0.3461 | -1.433 |
|  | Naming Order x Naming Condition | 0.0682 | 0.0521 | 1.308 |
|  | *Random effects* | Variance | Deviation |  |
|  | Participants |  |  |  |
|  | Intercept | 5.3898 | 2.322 |  |
|  | Naming Condition | 0.126 | 0.355 |  |
|  | Semantic categories |  |  |  |
|  | Intercept | 0.2266 | 0.476 |  |
|  | *Goodness of fit* |  |  |  |
|  | Log likelihood | -30877.6 |  |  |
|  | REML deviance | 61755.2 |  |  |

The maximal informative random structure for modeling participants’ naming latencies included random slopes for the factor naming condition for participants and random slopes for the factor naming order for semantic categories. As in Experiment 1, Naming Order and Repetition each contributed significantly to predicting naming latencies. Furthermore the model identified a significant interaction between Naming Order and the second to third repetition, which is probably caused by a spurious but rather prominent increase in naming latencies from Ordinal Position 1 to Position 2 in the third repetition cycle. Since this effect is not informative to the study’s hypotheses, and was also not found in Experiment 1, we will not discuss further this particular finding. Most importantly for the current study, our LMM model did not support the predicted interaction of Naming Order with Naming Condition. We conclude that also in Experiment 2 the partner’s naming task did not affect the participants’ own naming responses.

Electrophysiological response while participant names pictures. Experiment 2 replicates Experiment 1’s finding of a posterior positivity 250 to 400ms after participants initiated their naming response. The amplitude of this positivity increased in size from the first to the last picture named within a given semantic category, see Figure 4a. In line with our behavioral findings, there was no notable difference between the Joint Naming and the Single Naming condition.

**Figure 4.**
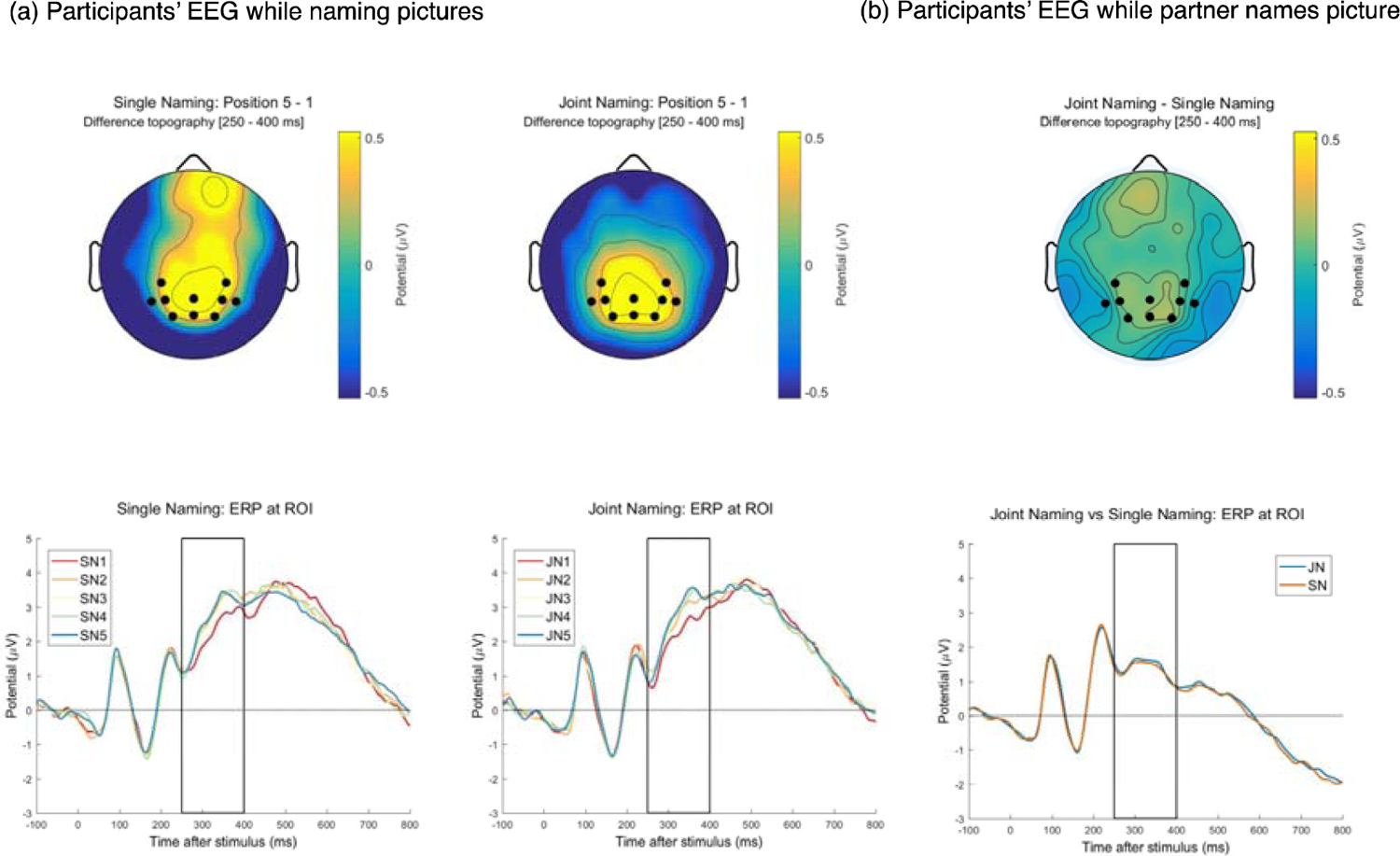
EEG results from Experiment 2. (a) Topographies (top) and waveforms (bottom) of event-related potentials elicited during participants’ presumed lexical access (250-400ms) when naming pictures of the fifth relative to the first ordinal position within a given semantic category for Single Naming (SN; right) and Joint Naming (JN; left) separately. (b) Topographies (top) and waveforms (bottom) of event-related potentials elicited in participants during trials in which they believe their partner is naming pictures (Joint Naming; JN) relative to trials in which pictures are presented only visually (Single Naming; SN) in the time-window of presumed lexical access (250-400ms). The electrodes of the region of interest are marked in black in the topography and were pooled to generate the waveforms. The time window of interest is marked by black box in the waveform. See the online article for the color version of this figure.

Correspondingly, our LMM model, with a random structure consisting of a random slope for naming conditions to model by-subject effects and a random intercept for semantic categories, yielded a significant contribution of the predictor naming order. None of our other predictors, Naming Condition and Repetition, contributed to our model.

A cluster-based permutation test searched for an effect of naming order by comparing EEG recordings during the first and last naming within a given semantic category, see Figure B, Appendix B. This analysis confirmed the increase in positivity in posterior electrodes reported above with a significant positive cluster (p = .00) reaching from about 275ms to 475ms and encompassing posterior as well as central electrodes. This cluster is quite comparable to the cluster found in Experiment 1. Furthermore, we found a significant negative cluster (p = .03) in the approximate time window 450-550ms encompassing mainly posterior and some lateral electrodes. Despite some smaller differences in involved electrodes and time window this cluster largely resembles the negative cluster found in Experiment 1.

Furthermore, this contrast yielded three additional significant clusters, which had not been found in Experiment 1: a positive cluster (p = .04) including frontal electrodes between about 200-250ms, a negative cluster (p = .03) including posterior electrodes in the time windows 200-250ms, and a negative cluster (p = .03) including left-lateralized electrodes in the time window 350-400ms.

As in Experiment 1, no significant clusters emerged from permutation tests searching for an effect of Naming Condition or for an interaction effect between Naming Order and Naming Condition. Electrophysiological response during partners’ lexical access. As in Experiment 1, we searched for evidence of a posterior positivity during those trials in which the task partner named pictures. However, such a positivity was not observed in either of our contrasts, (1) comparing the first and the last picture named within a given semantic category, (2) comparing the Joint Naming with the Single Naming condition, (3) comparing the difference between first and the last picture naming for each naming conditions., see Figure 4b. Accordingly, the specified LMM with a random slope for Naming Condition for subjects and a random intercept for semantic categories found no significant contribution of our predictors Naming Order, Naming Condition and Repetition. Please see Table 2 for model details.

Exploratory analyses (0-700ms). A cluster-based permutation test compared the participants’ EEG signal recorded while the task partner presumably named a picture (Joint Naming) to the EEG signal recorded while nobody named the presented picture (Single Naming). No significant clusters emerged from this comparison.

### Discussion

The findings of Experiment 2 resemble in remarkable accuracy those of Experiment 1. While we find clear evidence of cumulative semantic interference, both in the behavioral and electrophysiological data, we do not find evidence for partner-elicited semantic interference. Despite our efforts to emphasize the partner’s involvement in the task participants did not appear to represent their partner’s naming response. This was evidenced in participants’ EEG, which virtually showed no difference between trials in which the partner named the picture and trials in which the picture was presented only visually. Under neither of these conditions could we observe any signs that participants were engaging in lexical access.

To reconcile the current findings with our previous findings (Kuhlen & Abdel Rahman, 2017) we combined the data of these five experiments. This pooled dataset revealed robust empirical evidence for partner-elicited semantic interference and points towards a moderating role of the task partner’s physical co-presence.

## Pooled Analyses

### Methods

#### Data sets

Our pooled dataset integrated the original raw behavioral data of five experiments: three experiments that were reported in Kuhlen and Abdel Rahman (2017) and the presently reported two experiments. This amounted to 144 participants in total (107 women, 37 men) between the ages 18 to 36. See Table 3 for the number of participants for each contributing experiment.

#### Materials and Procedures

The stimuli were identical across experiments. In our 2017 experiments the entire set of stimuli were presented to each participants once. In the present two experiments, the same stimuli set was presented three times to each participant. This amounted to a different maximum number of observations an individual participant could contribute (see Table 3).

The procedures were in large parts identical across all five experiments. In two experiments (Kuhlen & Abdel Rahman 2017’s Experiment 1 and 3) participants and their task partner were physically co-present in the same experimental cabin during the joint naming task. In three experiments (Kuhlen & Abdel Rahman 2017’s Experiment 2, and the presently reported Experiment 1 and 2) participants and their task partner were seated in separate experimental cabins.

#### Analyses

Analog to the procedures described in Experiment 1 and 2, naming latencies were corrected for outliers and log-transformed. Latencies were then modeled by a linear mixed effect model including the predictors naming condition and naming order. Our initially run model included a full random structure with intercepts and slopes, as allowed by our experimental design, for participants and semantic categories. Furthermore we included an additional random term for experiments to control for experiment heterogeneity. To probe a possible moderating effect of the task partner’s physical co-presence we ran a second LMM model that included this binary variable as a third, contrast-coded predictor. A third LMM model looked at the influence of our predictors naming condition and naming order separately for those experiments with physically co-present and those with remote task partners.

## Results

### Partner-elicited interference

Over all datasets, participants named pictures with an average mean latency of 916ms (Joint Naming: 916ms; Single Naming: 916ms). Naming latencies increased over ordinal position by 13ms (cumulative semantic interference). Furthermore, this increase was steeper in the Joint Naming condition (16ms) than in the Single Naming condition (11ms), which yielded a significant interaction between naming order and naming condition (see Table 4 for model details and Figure 5 Panel A). This demonstrates a small, but statistically significant effect of partner-elicited interference over all five experiments. Our final model included random intercepts for subjects, semantic categories and experiments, and slopes for naming order within semantic categories and slopes for naming condition within experiments.

**Figure 5.**
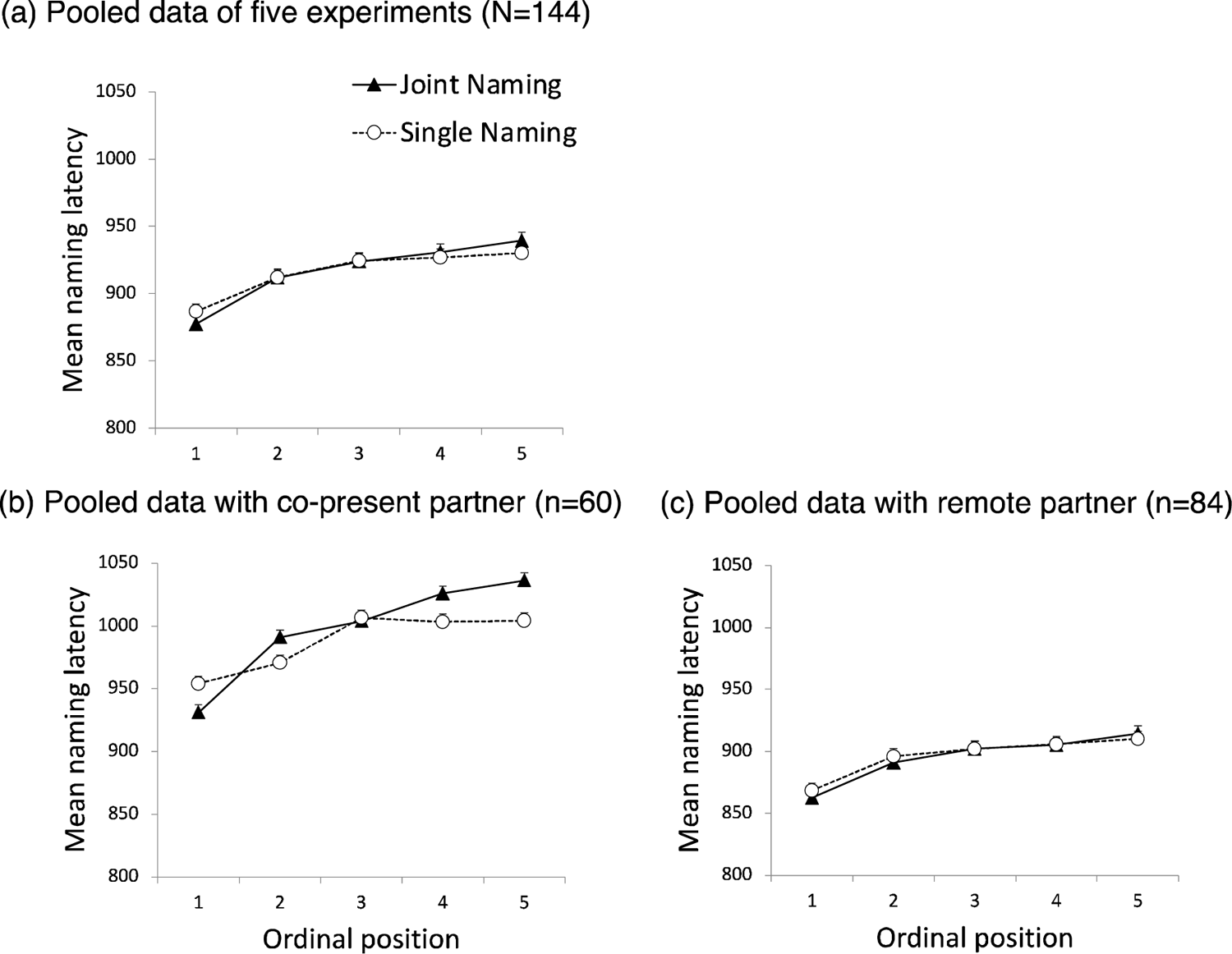
Data pooled from five experiments on partner-elicited cumulative semantic interference. Mean naming latency and standard error (in milliseconds) for each ordinal position and naming condition. (a) Results of all experiments. (b) Results of two experiments with physically co-present task partner. (c) Results of three experiments with remotely located task partner.

**Table 3:** Number of participants and experimental trials for experiments with physically co-present and remotely located task partners entering the pooled dataset.

| Data sets | Number of participants<br>(male, female) | Maximum number of<br>experimental trials<br>per participant | Location of task partner |
| --- | --- | --- | --- |
| Kuhlen & Abdel Rahman (2017), Exp. 1 | 24 (6, 18) | 160 | co-present |
| Kuhlen & Abdel Rahman (2017), Exp. 2 | 24 (2, 22) | 160 | remote |
| Kuhlen & Abdel Rahman (2017), Exp. 3 | 36 (9, 27) | 160 | co-present |
| Kuhlen & Abdel Rahman (current study), Exp. 1 | 30 (9, 21) | 480 | remote |
| Kuhlen & Abdel Rahman (current study), Exp. 2 | 30 (11, 19) | 480 | remote |

**Table 4:**
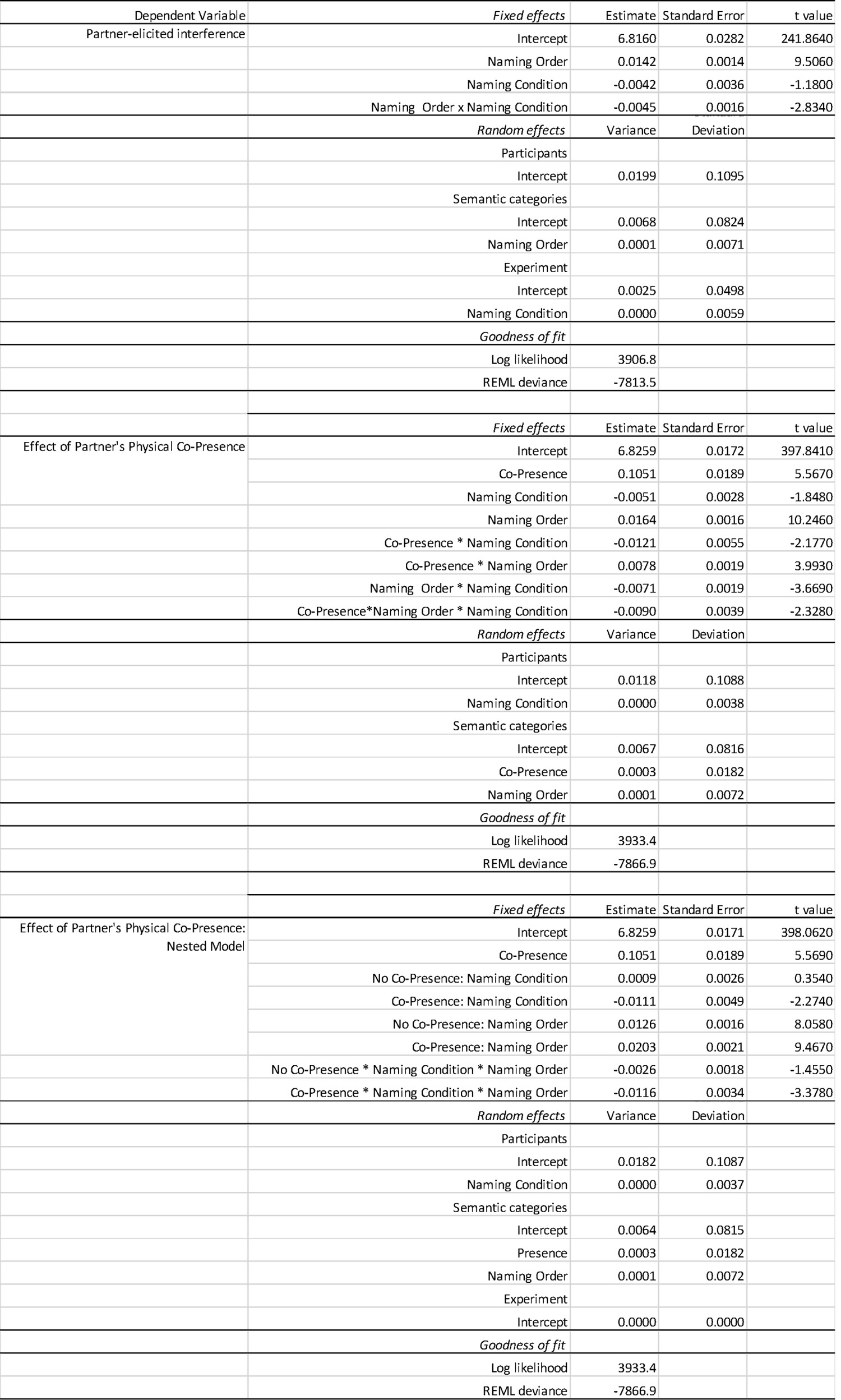
Fixed-effect estimates, standard errors, and t-values for the selected models of our pooled datasets; estimates of the variance and square root (standard deviations) of the random effect structure and goodness-of-fit statistics. Fixed effects are considered significant if |t| ≥ 1.96 (cf. Baayen, Davidson, & Bates, 2008).

Task partners’ physical co-presence. The task partner’s physical co-presence amplified partner-elicited interference. While, statistically, partner-elicited interference remained significant overall, speakers with physically co-present partners slowed down

14ms more in the Joint Naming compared to the Single Naming condition. Speakers with remote partners slowed down only 3ms, see Figure 5 Panel B and C. This yielded a significant three-way interaction in a model that included physical co-presence as predictor (see Table 4 for model output).

Furthermore, the partners’ physical presence appeared to slow down participants’ naming response overall, by about 100ms. As indicated by a significant interaction between the partner’s presence and naming condition this general slowing of naming latencies with physically present partners was particularly pronounced in the Joint Naming condition. Presumably, in this condition the general slowing due to the partner’s presence is amplified through the additional interference elicited by the partner’s naming responses. Lastly, our model revealed an interaction between the physical presence of a partner and naming order, indicating a stronger cumulative interference effect when partners were co-present (yet cumulative interference remains statistically significant in both settings, see also model reported next).

We explored further the pronounced partner-elicited interference with physically co-present compared to remotely located partners through a third, nested model. This model supports the conclusion that partner-elicited semantic interference is statistically significant only in settings in which the task partner is physically present, see Table 4 for model output.

## Discussion

Our pooled data analyses demonstrate, once more, robust cumulative semantic interference. What is more, over all five experiments, we find empirical evidence for partner-elicited interference. This effect is amplified in a setting in which the task partner is physically co-present. Note that this setting does not necessarily entail that participants were able to hear their partner’s verbal response. In one of two experiments with a physically co-present partner the partner’s response was masked (but not in the other). We conclude that, despite the absence of partner-elicited interference in the presently reported Experiment 1 and 2, partner-elicited interference is real. The reason why we did not find partner-elicited interference in the current study may lie in the generally small size of the effect and in our setting, which physically separated our task partners. We will discuss the implications of these results in the following section.

### General discussion

When naming pictures, speakers show increased naming latencies with each additional picture they name within the same semantic category (cumulative semantic interference; see e.g., Belke, 2013; Costa et al., 2009; Howard et al., 2006; Navarrete et al., 2010). Previous studies have demonstrated that naming latency not only increases in response to participants’ own prior naming of within-category pictures, but also in response to a task partner naming the pictures (Hoedemaker et al., 2017; Kuhlen & Abdel Rahman, 2017). In this study we used electroencephalographic recordings (EEG) to gain insights into the mechanisms underlying this partner-elicited semantic interference. It has been suggested that speakers predict their conversational partner’s utterances by simulating the cognitive processes involved in producing it (Pickering & Garrod, 2013). We therefore proposed that partner-elicited semantic interference is caused by participants engaging in lexical access for pictures that are to be named by their task partner. To investigate this proposal, we focused on a particular event-related potential, a posterior positivity 250-400ms after picture onset, as a proxy for (simulated) lexical access. Specifically, we compared trials in which the partner named pictures to those trials in which pictures were presented only visually (but named by nobody). Our experiments were pre-registered (Kuhlen & Abdel Rahman, 2017). Because Experiment 1 did not yield the expected results we ran a second experiment with slightly modified procedures. Experiment 2 replicated almost exactly the results of Experiment 1. We therefore combine the discussion of both experiments.

Both experiments show consistent evidence, in participants’ behavioral responses as well as in their EEG signal, for cumulative semantic interference: Naming latencies increased with each picture named within a given semantic category, replicating once more this quite robust effect (e.g., Belke, 2013; Costa et al., 2009; Howard et al., 2006; Navarrete et al., 2010). Paralleling this behavioral finding, a posterior positivity could be observed within the electrode positions and the time window reported in previous studies (precisely: Rose & Abdel Rahman, 2016; overlapping: Costa et al., 2009). Since the time window of the component aligns with the presumed time window of lexical access (Indefrey, 2011; Indefrey & Levelt, 2004) and gains in amplitude corresponding to the behavioral interference effect, it is commonly interpreted as reflecting the ease with which lexical access can be achieved (Costa et al., 2009). Also in our study the component increased in amplitude from the first to the last ordinal position. Both findings, increasing naming latencies and increasing positivity, indicate that participants experienced interference when naming pictures within the same semantic category. This finding is consistent with the proposal that semantically related lexical entries compete with each other for lexical selection.

However, our hypotheses concerning partner-elicited semantic interference were not supported by the data. Based on previous studies (Hoedemaker et al., 2017; Kuhlen & Abdel Rahman, 2017) we had expected that participants would seek lexical access for pictures named by their task partner. This would produce additional lexical competitors and would thus result in overall stronger interference for those semantic categories named together with the task partner (vs. those named exclusively by the participant). However, in both conditions naming latencies increased at a comparable rate. Since the overall increase in latencies was rather small compared to previous reports, the most likely explanation for this data pattern is that participants did not seek lexical access for pictures they did not name – regardless of whether these were named by the partner or were presented visually.

Correspondingly, we did not find any sign of lexical access for pictures named by the partner in participants’ EEG. This is evidenced by the absence of the posterior positivity we observed during participants’ own picture naming in trials in which the task partner named pictures. It has been proposed that in joint actions representations of the partner’s task need to be eventually inhibited in order to avert execution (Sebanz et al., 2006). Based on this account, one possible explanation for the lack of lexical access in our experiments could be that the partner’s lexical response was not represented to the level of lexical access. Instead, the simulation of the partner’s language production may have been inhibited at an earlier stage. However, we did not find any signs of increased inhibitory processes in trials in which task partners named pictures. In fact, exploratory analyses, including all electrodes and a large time window of 100-700ms, found no significant differences between trials in which the partner named the picture and trials in which the pictures were presented only visually. We therefore conclude that participants did not represent at all their partner’s naming response.

Our prior power analysis and the replication of these findings over two experiments make a spurious null effect rather unlikely. Yet, the present experiments stand in contrast to previous studies that report partner-elicited semantic interference in nearly identical (Experiment 2, Kuhlen & Abdel Rahman, 2017) or comparable (Experiment 1 and 2, Kuhlen & Abdel Rahman, 2017; Hoedemaker et al., 2017) settings. Our present findings are also in contrast to studies employing other types of joint-picture naming tasks (Baus et al., 2014) and to a large body of literature on joint action in task domains that do not involve language processing (for overview see Knoblich, Butterfill, & Sebanz, 2011). To reconcile these findings we integrated data from five experiments on partner-elicited interference. Over all these experiments, partner-elicited interference emerged as a small, but significant effect.

One explanation for the lack of representing the partner (or their response) in the current two experiments is that the two task partners were spatially separated during the picture naming. Indeed, our pooled analyses suggest that settings with remotely located task partners elicit substantially less, or even no, interference through the partner’s naming response. A physical separation of task partners may undermine the social nature of task. Indeed, in both experiments reported in this manuscript, a number of participants had to be excluded (and replaced) because they indicated doubt that the partner was indeed naming the pictures. And even among the remaining participants a high number of participants reported that they did not feel the setting was a joint action and they indicated that it had made no difference to them whether the partner or nobody named the picture.

The majority of studies investigating joint action employ settings in which the task partners are physically co-present. Nevertheless, there are studies that report partner co-representation of remotely located task partners (e.g., Gambi et al., 2015; Ramnani & Miall, 2004; Tsai et al., 2008). In their review of joint action literature Knoblich and colleagues (2011) speculate that constant feedback from the task partner may be necessary to maintain a representation of a remotely located task partner (but see Vlainic et al., 2010). In our setting feedback from the partner was provided in form of mock naming latencies (which could be inferred by the speed at which pictures disappeared in the partner trials). Yet this form of feedback may not have been salient enough to maintain a stable partner representation. The circumstances under which task partners represent each other’s actions (and to what level of detail) is thus an important question to be addressed in future research. The outcome of the present experiments reminds us once more that when transferring social interaction into an experimentally controlled setting great care needs to be taken that the essential features of social interaction are not lost (for a related argument see Kuhlen & Brennan, 2013).

Our failure to observe partner-elicited interference with remotely located task partners may be connected to our finding that semantic interference in general seems reduced in these settings. In contrast, semantic interference was amplified when participants named pictures alongside a physically present partner. Furthermore, in settings with a physically present partner, we observed a pronounced slowing of overall naming latencies. Possibly, the experience of semantic interference, as well as the experience of partner-elicited semantic interference is tied to overall slower naming latencies. Indeed, distributional characteristics of naming latencies have been investigated in the context of semantic interference effects, and some of these studies have proposed that semantic interference is more pronounced in slower naming latencies (e.g., Scaltritti et al., 2015; but see Roelofs & Piai, 2017). Yet, this literature investigates semantic interference elicited in picture-word interference paradigms and ties the speed of the naming response to the processing of the distractor word manipulating the semantic context. In the context of continuous naming tasks, as we employ in the present experiments, we are not aware of a connection between general naming latencies and the size of the semantic interference effect. This might be an interesting avenue for future research.

There is a growing number of studies employing joint picture naming tasks that report only limited, or no, representation of the partner’s naming response (Brehm et al., 2019; Gambi, Van de Cavey, & Pickering, 2015; Hoedemaker & Meyer, 2019). These studies come to the conclusion that speakers may represent their partner’s naming response, but not necessarily to the degree that they seek lexical access for the pictures the partner names. This conclusion is supported by a body of literature investigating non-linguistic joint actions (for overview see Dolk et al., 2014). Most notably, in the so-called Social Simon task (Sebanz, Knoblich, & Prinz, 2003) two task partners each perform one part of a spatial compatibility task (e.g., participant: press left key for red stimuli; partner: press right key for green stimuli). When working together with a partner (vs. performing their part of the task alone), participants show a classic spatial compatibility effect that results from representing two stimulus-response alternatives. Participants slow down when the irrelevant location of the stimulus and the location of the response do not overlap (e.g., red stimuli presented on the right side requiring a left key response) – even though participants only have responsibility for one action. This suggests that participants represent their partner’s actions functionally similar to their own, which aligns with our initial hypotheses. However, an alternative explanation for this finding is that participants do not represent specific details of their partner’s task, but merely who needs to respond (e.g., participant: red; partner: green; Dolk et al., 2011; Philipp & Prinz, 2010; Vlainic, Liepelt, Colzato, Prinz, & Hommel, 2010; Wenke et al., 2011). This proposal is in line with the recent findings from joint picture naming that only find evidence for limited representations of the partner’s naming response (Gambi et al., 2015; Hoedemaker & Meyer, 2019). Yet the pattern of results found in our present experiments do not lend themselves to distinguishing between these two accounts. Since we found no indication that the partner’s actions were represented at all we cannot say anything about the nature of this representation. Our data pattern appears to suggest that the partner’s task was irrelevant and may not have registered with participants.

Despite our somewhat inconclusive findings regarding partner-elicited interference our study contributes to the growing body of literature investigating overt speech production with electroencephalographic recordings. Studies recording EEG during speech production are still rather rare due to the artifacts produced in the EEG signal by the muscle movements associated with speaking (Ganushchak, Christoffels, & Schiller, 2011). Different methods have been developed to address this challenge (e.g., Ouyang et al., 2016; Porcaro, Medaglia, & Krott, 2015; Vos et al., 2010). Based on these efforts consistent patterns of activation have been identified (e.g., Aristei, Melinger, & Abdel Rahman, 2011; Indefrey, 2011; Laganaro, 2014; Strijkers & Costa, 2016). One of them is the posterior positivity 250-400ms after picture onset, which we have targeted in this study. This component has been consistently observed in response to naming semantically related pictures in the continuous picture-naming paradigm and has been related to lexical selection (Costa et al., 2009; Rose & Abdel Rahman, 2016; for a similar posterior positivity related to lexical selection in the picture-word interference task, see Rose, Aristei, Melinger, & Abdel Rahman, 2019).

These studies also report the modulation of a second posterior component, a relative negativity around 400ms after picture onset, commonly labeled as N400 (Kutas & Federmeier, 2011). In the current two experiments we also observed such a relative negativity at posterior electrode sites. Comparable to previous studies the negativity we found was most pronounced at posterior electrode sites, at a time window of 450-600ms, and increased in negativity from the first to the last picture named within a semantic category. In studies on language comprehension the N400 has consistently been associated with semantic processing (for review, see e.g. Kutas & Federmeier, 2011; Lau, Phillips, & Poeppel, 2008; Rabovsky & McRae, 2014). Typically, the N400 has a large amplitude when processing semantic violations, for example when a word does not fit into the given sentence context. The N400 is also modulated by semantic properties associated with words: Amplitudes are smaller in response to words that are paired with semantically related compared to unrelated words (Bentin, Mccarthy, & Wood, 1985). And amplitudes are higher in response to processing words with many compared to few semantic features (Holcomb, Kounios, Anderson, & West, 1999; Kounios & Holcomb, 1994; Laszlo & Federmeier, 2011; Müller, Duñabeitia, & Carreiras, 2010; Rabovsky, Sommer, & Abdel Rahman, 2012b, 2012a).

In language production, however, the N400 is less extensively explored. In picture-word interference paradigms, a decrease in its amplitude was observed when naming target pictures after the presentation of a distractor word that corresponded to the picture name (Blackford, Holcomb, Grainger, & Kuperberg, 2012; Chauncey, Holcomb, & Grainger, 2009) or when naming pictures after a morphologically similar distractor word (Koester & Schiller, 2008). Most interesting in the current context, the N400 appears to be diminished when the distractor word is semantically related to the target picture (Blackford et al., 2012). Based on this finding the N400 has therefore been suggested to reflect semantic priming (see also Roelofs, Piai, Garrido Rodriguez, & Chwilla, 2016). In the current experiments we also find a relationship between the observed negative modulation and semantic context, but in the opposite direction: We observe an increase in negativity that corresponds to the increase in semantic interference observed in speaker’s naming latency. A similar finding is sometimes related to cognitive control and the resolution of lexical competition (e.g., Liotti, Woldorff, Perez, & Mayberg, 2000; Piai, Roelofs, & van der Meij, 2012; Rose & Abdel Rahman, 2016). Further studies are needed to understand the N400 component during language production and the role semantic context may play in its modulation.

In conclusion, the present two EEG experiments contribute to a growing number of experiments investigating language production in a social interaction. We investigated the question whether a task partner’s naming response is simulated to the degree of seeking lexical access on behalf of the partner. In contrast to previous studies, we find no evidence that a partner’s naming response is represented. A pooled analysis, including the presently reported two experiments and three previously published experiments on partner-elicited interference, suggests that our setting, which physically separated task partners from each other, may have affected whether, or to what degree, the partner’s response is represented. The outcome of the present experiments cautions to pay close attention to participants’ evaluation of the setting as a joint action. In addition, the experiments contribute to our understanding of the temporal dynamics underlying language production by providing converging evidence for stable electrophysiological markers associated with semantic interference.

## Acknowledgements

This research was supported by the grants KU3236/3 and AB277/11 from the German Research Council. We thank Anne Schmidtke, Anna Bakhireva, Joanna Rickman, Sophie Taylor, and Sonia Lyn for their help in the data collection, Antje Lorenz for discussion, and Guido Kiecker for technical support.

## Appendix A

**Figure A.**
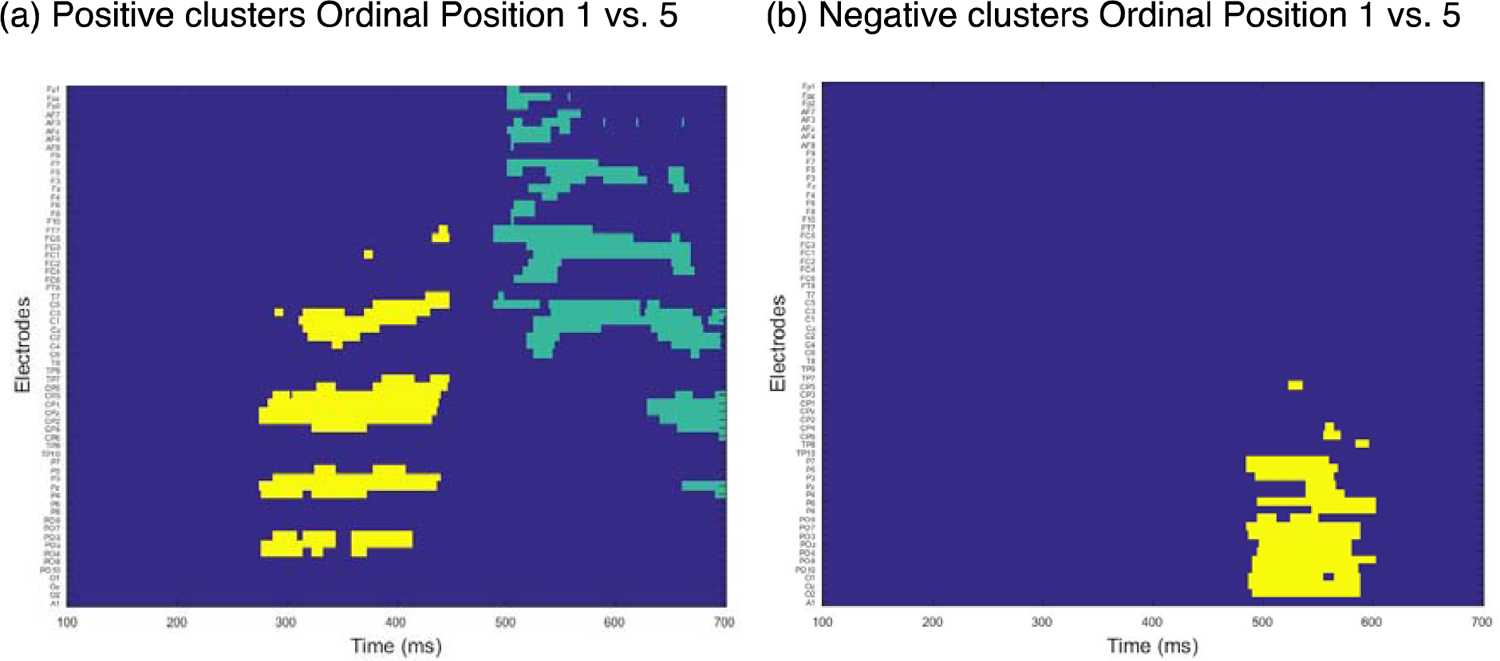
Results of permutation test from Experiment 1 on trials in which participant names pictures over all electrodes and the time window 100-700ms after pictures onset. Colors indicate individual clusters. (a) Positive clusters comparing ordinal position 1 with ordinal position 5. (b) Negative clusters comparing ordinal position 1 with ordinal position 5.

## Appendix B

**Figure B.**
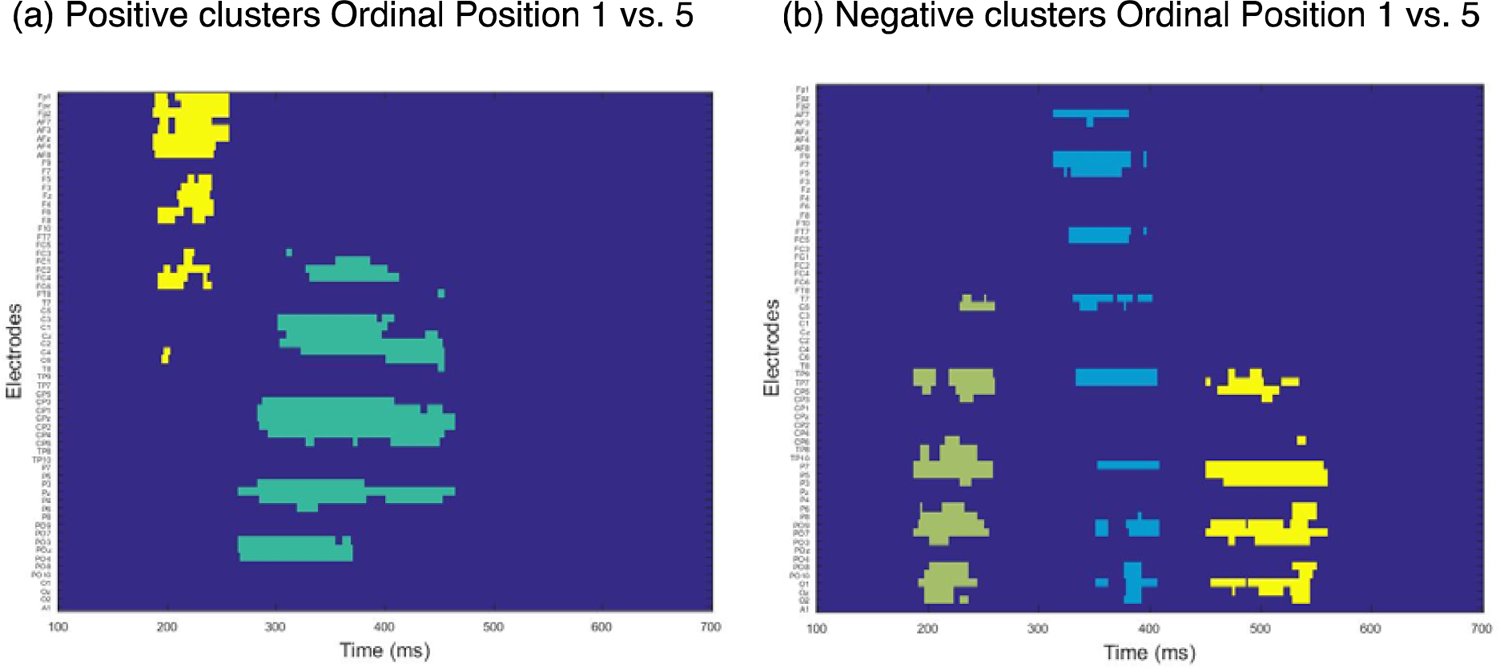
Results of permutation test from Experiment 2 on trials in which participant names pictures over all electrodes and the time window 100-700ms after pictures onset. Colors indicate individual clusters. (a) Positive clusters comparing ordinal position 1 with ordinal position 5. (b) Negative clusters comparing ordinal position 1 with ordinal position 5.

## Footnotes

A random structure modeling individual items does not significantly change the effect of our main predictors.

